# Size uniformity of animal cells is actively maintained by a p38 MAPK-dependent regulation of G1-length

**DOI:** 10.1101/119867

**Authors:** Shixuan Liu, Miriam B. Ginzberg, Nish Patel, Marc Hild, Bosco Leung, Yen-Chi Chen, Zhengda Li, Nancy Chang, Shulamit Diena, Yuan Wang, William Trimble, Larry Wasserman, Jeremy Jenkins, W. Kirschner Marc, Ran Kafri

## Abstract

Animal cells within a tissue typically display a striking regularity in their size. To date, the molecular mechanisms that control this uniformity are still unknown. We have previously shown that size uniformity in animal cells is promoted, in part, by size-dependent regulation of G1 length. To identify the molecular mechanisms underlying this process, we performed a large-scale small molecule screen and found that the p38 MAPK pathway is involved in coordinating cell size and cell cycle progression. Small cells display higher p38 activity and spend more time in G1 than larger cells. Inhibition of p38 MAPK leads to loss of the compensatory G1 length extension in small cells, resulting in faster proliferation, smaller cell size and increased size heterogeneity. We propose a model wherein the p38 pathway responds to changes in cell size and regulates G1 exit accordingly, to increase cell size uniformity.

**One-sentence summary:** The p38 MAP kinase pathway coordinates cell growth and cell cycle progression by lengthening G1 in small cells, allowing them more time to grow before their next division.

## Introduction

Homogeneity of cell size is a characteristic feature of healthy tissues. While cells from different tissues vary widely in size, within a given tissue cells are strikingly similar in size (1). In a separate study (co-submitted), we showed that size uniformity in populations of animal cells results from two separate processes. Firstly, during G1, a size-dependent process delays S-phase entry in small cells, giving them more time to grow before their next division. Additionally, twice during the cell cycle, cellular growth rates are adjusted so that small cells actually grow faster than large cells. The molecular mechanisms underlying these regulatory behaviors have not yet been identified. In yeast, the existence of a cell size checkpoint has long been postulated (2, 3). However, the molecular mechanisms maintaining that checkpoint also remain elusive.

The mammalian p38 MAPK pathway participates in numerous biological processes, including the regulation of cell cycle checkpoints. In response to DNA damage or oxidative stress, p38 is activated and induces a cell cycle arrest (4, 5). Hyperosmotic conditions that shrink cell volume also strongly activate p38 (6–8). Furthermore, upregulation of the p38 MAPK pathway can lead to increased cell size (9–13). These observations raise the possibility that p38 may function to regulate cell size checkpoints in animal cells.

## Results

To identify the molecular pathways linking cell size and cell cycle progression, we searched specifically for perturbations that could disrupt this link. We screened two compound libraries, known as the Novartis MOA (Mechanism-of-Action) box and Kinome box, which together included over 3,000 compounds. The MOA Box contains a dynamically-managed annotated list of compounds curated to maximize coverage of targets, pathways, and bioactivity space (Figure 1 -Figure supplement 1), to facilitate biological discovery by screening and profiling experiments. The Kinome Box is a library containing a wide range of kinase inhibitors that were selected based on their efficiency and specificity (primary targets IC50<1 μM, <25 total targets). Compounds from both libraries are thoroughly annotated for primary targets, off-target effects and corresponding potencies.

To initiate the screen, unsynchronized HeLa cells were treated with compounds for 24 hours, after which cells were fixed and both the size and cell cycle stage of each cell was measured. The cells expressed the FUCCI cell cycle reporters mAG-hGem and mKO2-hCdt1 (14), enabling sensitive detection of G1 progression and exit. Additionally, to assay progression through S phase, fixed cells were stained with DAPI. To measure cell size, cells were also stained with a protein dye that provides an accurate measure of cell mass (15). Following drug treatment, samples were imaged by fluorescence microscopy, and an automated image-processing pipeline was developed to quantify fluorescence intensity within each cell. With this screening strategy, both cell size and cell cycle stage of each cell was measured. Each drug treatment yielded a multi-dimensional joint distribution of cell size and cell cycle indicators (Figure 1A). After subsequent data analysis, the single cell measurements from each treatment can be classified into discrete cell cycle stages (Figure 1 -Figure supplement 2). Cell count, average cell size, and cell size variability were also calculated, to evaluate how each compound influences cell proliferation and size regulation.

**Figure 1.**
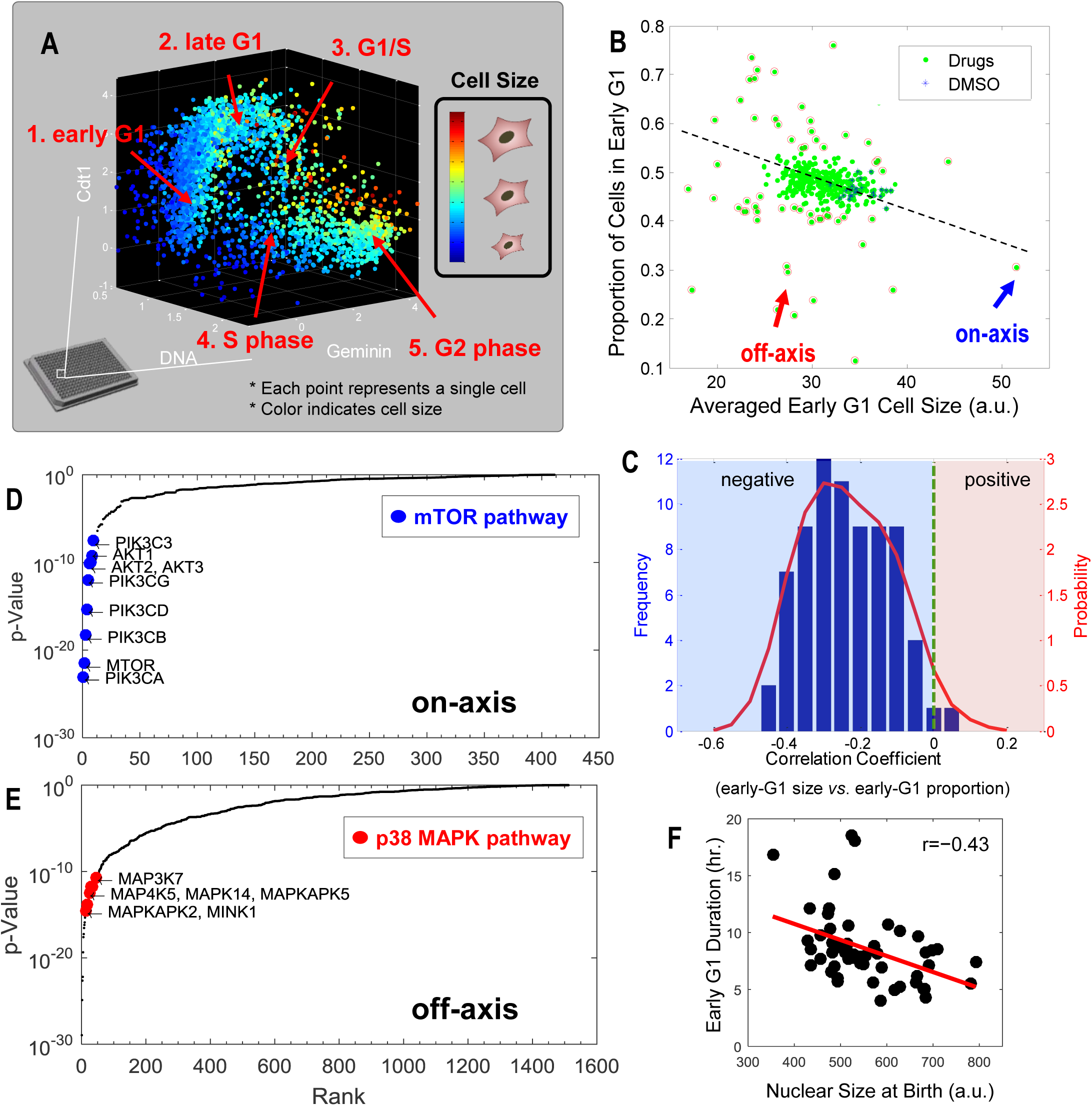
Results from a small molecule screen implicate the p38 MAPK pathway in the coordination of cell size and progression through G1. **A**. Scatter plot showing the relationship of cell size and cell cycle stage in one example screened condition. Single cell data is classified into discrete cell cycle stages according to the 3 cell cycle indicators; Geminin, Cdt1 and DNA (**Figure 1 -Figure supplement 2**). **B**. Average size of early G1 cells is negatively correlated with the fraction of cells in early G1. The scatterplot displays the result from one example 384-well plate. Each data point represents one particular compound treatment (or DMSO control). Red circles highlight the conditions that significantly affect the size of early G1 cells or the proportion of cells in G1. The arrows designate examples of on-axis and off-axis compounds (also see **Figure 1 -Figure supplement 3**). **C.** Distribution of correlation coefficients between size of early G1 cells and the fraction of cells in G1, calculated for all screened plates (as described in Methods), demonstrating that the two variables are significantly negatively correlated (*p*<10^-16^). **D, E**. Ranked *p*-values from the target enrichment analysis of on-axis and off-axis compounds respectively (Fisher’s exact test). Components of the mTOR pathway and p38 MAPK pathway are among the top-ranked hits that are highlighted. **F**. Live-cell imaging confirms that the duration of early G1 is negatively correlated with cell size at birth (N=53). The results shown here are representative of three replicate experiments.

In an unsynchronized population, the proportion of cells in a particular cell cycle stage reflects the duration of that stage relative to the entire cell cycle (15). Plotting the proportion of cells in early G1 versus the average size of early G1 cells, in each screened condition, consistently revealed a negative correlation between these quantities (Figure 1, B and C). Treatments that reduced the size of cells in early G1 typically yielded larger proportions of cells in G1, suggesting that G1 duration depends on a cell’s initial size. Live-cell tracking of single cells with time-lapse microscopy confirmed that there is a negative correlation between the nuclear size (a correlate of cell size (manuscript co-submitted)) of a cell at birth and its G1 duration (Figure 1F), further supporting the hypothesis that G1 length is regulated in a size-dependent manner to promote uniformity among cells.

To investigate the pathways that link cell size to G1 length, we identified compounds that altered the average cell size, fraction of cells in G1, or both (*i.e*. outliers on the plot in Figure 1B and Figure 1 -Figure supplement 3). We sorted these “hits” into two different types. The first, which we call *on-axis compounds*, perturbed both cell size and the proportion of cells in G1, but maintained the coordination between the two. In other words, on-axis compounds produced reciprocally correlated influences on cell size and the fraction of cells in G1. The second class of hits, *off-axis compounds*, are compounds that disproportionally affected either size or cell cycle progression, resulting in paired values of cell size and G1 fraction that lie off-axis to the trend defined by the majority of screened conditions in Figure 1B. These off-axis compounds may reveal a disruption of the coordination of size and G1 length.

To identify the pathways that are associated with either the off-axis or on-axis phenotypes, we used the compound annotations to perform target-enrichment analysis with Fisher’s exact test. Specifically, we asked for proteins that are significantly overrepresented in the combined list of targets of either the on-axis or off-axis compounds. This analysis avoids the risk of formulating a hypothesis based on a single drug. Interestingly, we found that the proteins most frequently targeted by the on-axis compounds are PI3K, Akt and mTOR (Figure 1D). This suggests that while the PI3K/Akt/mTOR pathways regulate cell size, these pathways are not involved in the coordination of cell growth and cell cycle progression.

By contrast, the MAP Kinases were highly enriched among the targets of off-axis compounds (Figure 1E). Notably, multiple components of the p38 MAPK pathway, including p38α (MAPK14), MK2 (MAPKAPK2, a direct substrate of p38) and TAK1 (MAP3K7, an essential upstream activator of p38) are ranked in the top 1% of significant targets (*p*-value<0.01). Over 35 compounds that target the p38 pathway components highlighted in Figure 1E, consistently displayed an off-axis phenotype, suggesting the involvement of the p38 MAPK pathway in the coordination of G1 length with cell size. As an additional test for pathways that control cell size, we ranked compounds based on their effect on cell size variability. Using a similar target-enrichment approach, we generated a list of candidate proteins whose inhibition is associated with increased variability in size. MK2, a downstream substrate of p38 was the highest-ranking target in this list (Figure 1 -Figure supplement 4).

To test whether the p38 MAPK pathway coordinates G1 length with cell size, we developed a robust assay to quantify this coordination in non-cancerous RPE1 cells. The mTOR inhibitor, rapamycin, is an on-axis compound that perturbs cell size but maintains the coordination of size with G1 length. In a different study (co-submitted), we showed that rapamycin produces a 60% decrease in the average rate of cell growth which is partially counteracted by an 80% increase in cell cycle length to limit the drug's effect on cell size. As shown in Figure 2A, higher concentrations of rapamycin correspond to smaller G1 cells and longer G1 duration, producing a robust negative correlation. To test if p38 is required for the negative correlation of cell size with G1 length, we examined a panel of specific inhibitors against MAPK pathways and tested whether any of these inhibitors break the negative correlation induced by rapamycin. Significantly, inhibition of p38, but not the JNK or ERK pathways, consistently breaks the negative correlation between size and proportion of cells in G1 (Figure 2 B-E, Figure 2 -Figure supplement 1 and Figure 2 -Figure supplement 2). This result was observed with multiple chemical inhibitors of p38, strongly implicating the p38 pathway in the coordination of G1 length with cell size.

**Figure 2.**
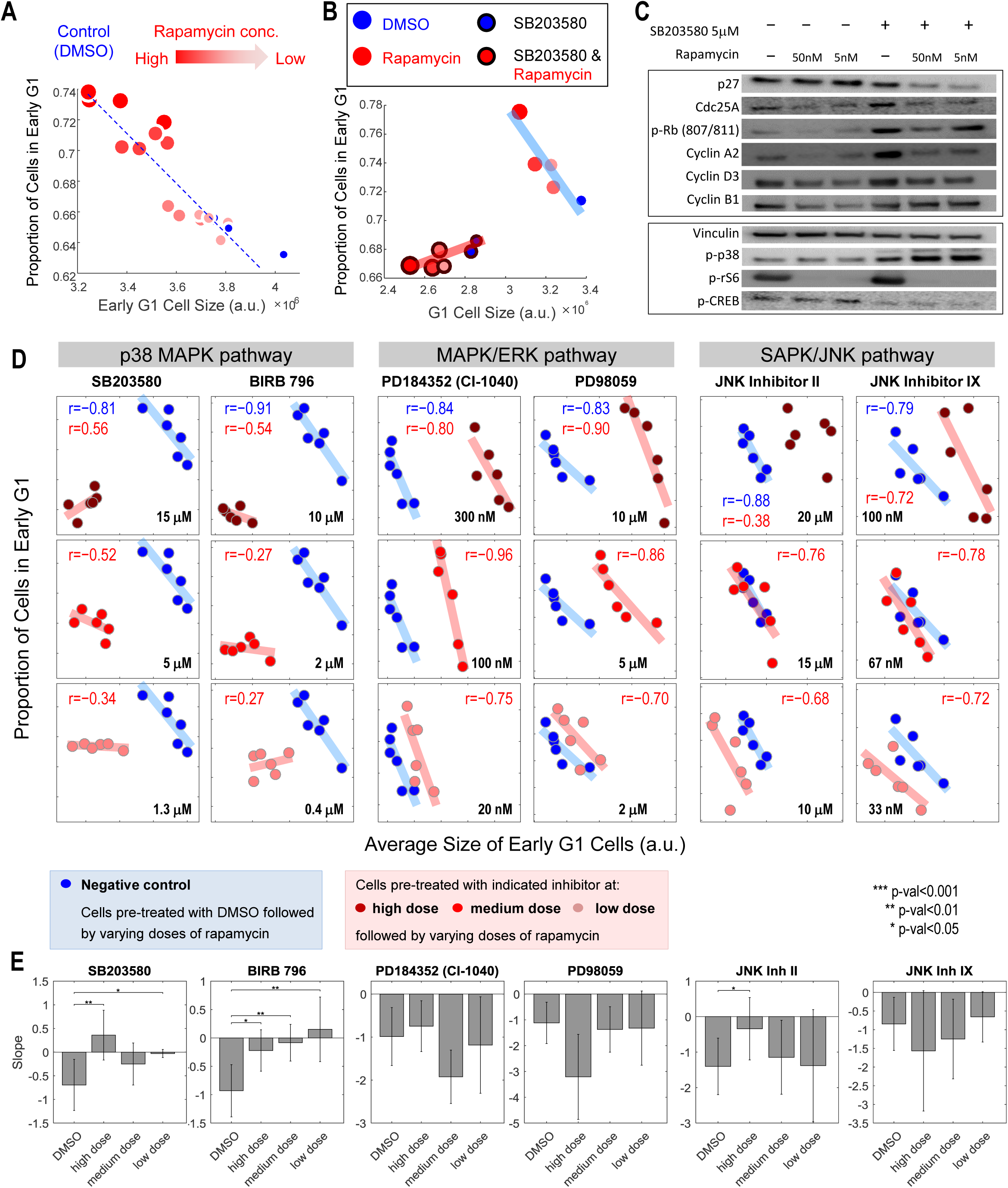
Pharmacological inhibition of the p38 MAPK pathway results in a loss of coordination of cell size with G1 length. **A**. Cells treated with varying concentrations of Rapamycin (mTOR inhibitor) demonstrate a robust negative correlation between the average size of early G1 cells and the proportion of cells in early G1. **B**. The negative correlation between cell size and proportion of cells in G1 disappears when cells are pre-treated with a specific inhibitor of p38 MAPK (5 μM SB203580). **C**. Westem-blot of whole cell lysates from the conditions shown in **Figure 2B. D**. Inhibition of the p38 MAPK pathway, but not the MAPK/ERK or SAPK/JNK pathways, disrupts the correlation between cell size during early G1 and the proportion of cells in G1. Each data point was measured from a cell population with a minimum of 7,000 cells. The results shown here are representative of three independent experiments. **E**. The fitted slope of measurements shown in **Figure 2D**. For each compound treatment, its fitted slope is compared with the slope of the control (DMSO) from the same experiment. Significance was calculated with one-tailed Student *t*-test (H_0_: slope_drug_<=slope_control_).

Knockdown of p38γ and p38δ, by siRNA drastically disturbs the negative correlation between cell size and proportion of cells in G1, while knockdown of p38α and p38β only partially weakens the correlation (Figure 3 A and B, Figure 3 -Figure supplement 1). This result strongly suggests functional divergence among the p38 isoforms with regard to size control (13, 16). One difference between the chemical inhibition and the knockdown experiments is a greater reduction in cell size observed following chemical inhibition of p38. A possible explanation for this difference is the lower specificity of chemical inhibitors as compared to genetic perturbations. Functional redundancy between different p38 isoforms, for example, can partially compensate for the siRNA but not for the pharmacological perturbations. Alternatively, it may be that transfection with siRNA activates nonspecific cellular responses, such as upregulation of innate immune pathways, that lead to an increase in size (17–19). Nonetheless, these results collectively demonstrate that both chemical and genetic inhibition of p38 activity result in loss of coordination between cell size and G1 length.

**Figure 3.**
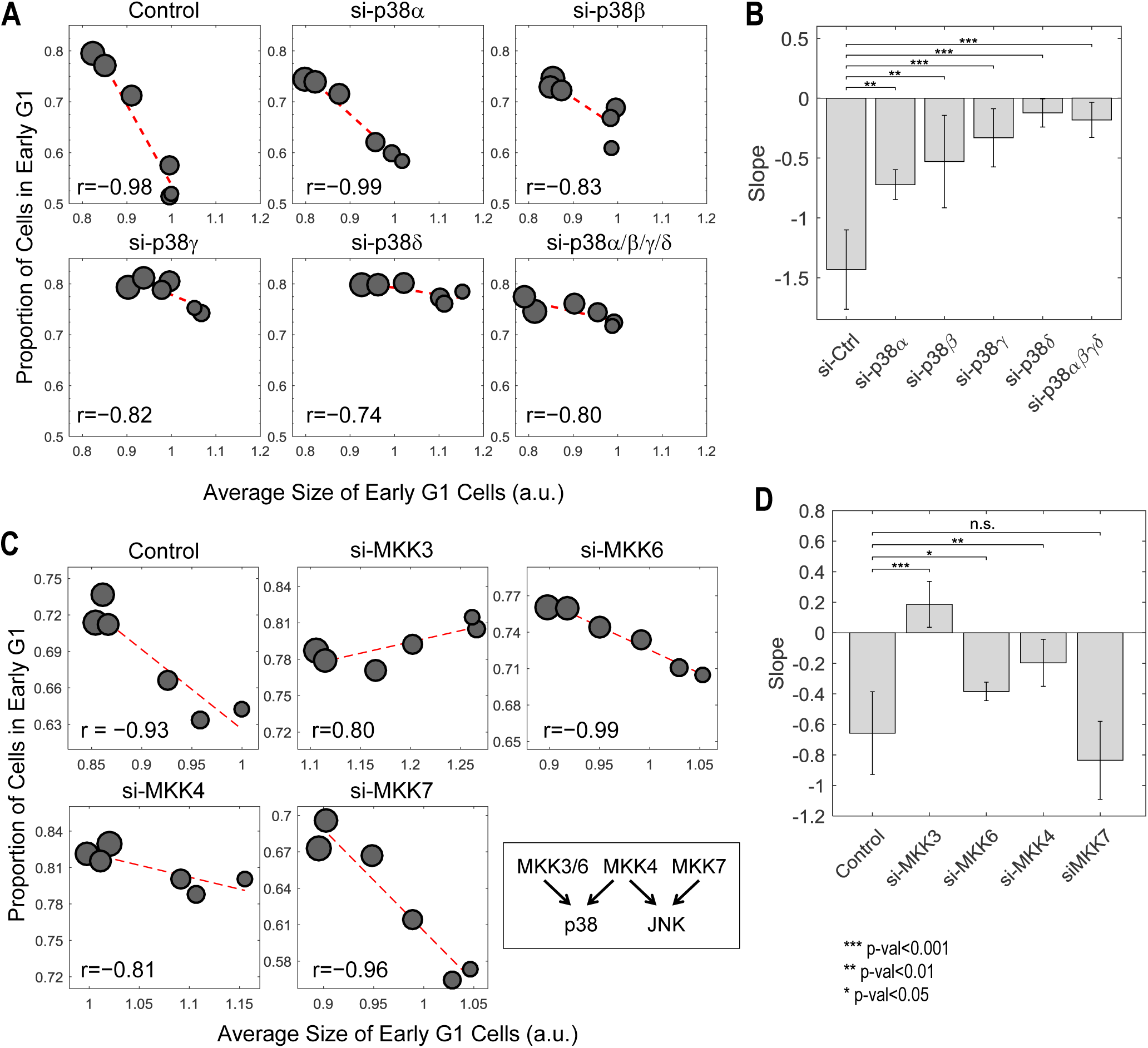
Knockdown of p38 pathway disturbs the negative correlation between cell size and proportion of cells in G1. **A**. Knocking down p38α/β partially weakens the negative correlation between size and proportion of cells in G1, while knockdown of p38γ/δ drastically disturbs the correlation. **C**. The negative correlation between cell size and proportion of cells in G1 is disturbed when cells are transfected with siRNA against MKK3/4/6 but not MKK7. Each data point in **Figure 3A and C** is measured from a cell population with a minimum of 3000 cells. The results shown in **Figure 3A and C** are representative of two/three independent experiments with duplicates or triplicates. **B, D**. The fitted slope of results shown in **Figure 3A and C**. All error bars indicate 90% confidence bounds. Analysis is performed with the same method as indicated in Fig 2.

To further test the relationship of p38 to size control, we perturbed MKK3/6/4 by siRNA, which are upstream activators of the p38 pathway. As a control, we also knocked down MKK7, an upstream regulator of JNK. Consistent with results of the chemical inhibitors, knocking down MKK3/6/4 disturbs the negative correlation between cell size and proportion of cells in G1 while knockdown of MKK7 leave the coordination of G1 and cell size intact (Figure 3 C and D, Figure 3 -Figure supplement 1).

The p38 MAPK pathway participates in cell cycle checkpoints by phosphorylating cell cycle regulators including p27, cyclin D, Rb and CDC25 (4, 20–22). To test the mechanisms of the p38-mediated cell size checkpoint, we treated cells with rapamycin, an on-axis drug, to induce a size-dependent lengthening of G1. As expected, rapamycin treatment was accompanied by downregulation of positive G1/S regulators (*e.g*., Cyclin D, p-Rb, Cdc25A) and upregulation of negative regulators of G1 progression (*e*.*g*., p27^Kip1^) (Figure 2C). In contrast, inhibition of p38 disrupted the effect of rapamycin on these four cell cycle regulators (Figure 2C), resulting in the loss of compensatory G1 lengthening in small cells. These results suggest that the p38 pathway functions downstream of a cell size sensor to prolong G1 length in small-sized cells and promote cell size uniformity.

The hypothesis that p38 regulates cell-size-dependent cell cycle progression predicts that the p38 MAPK pathway is selectively activated in small cells. Figure 2C shows that p38 is, indeed, activated when cell size is decreased by inhibition of mTOR. While this result is consistent with size-dependent p38 activation, it could also have alternative explanations. p38 may be directly activated by the inhibition of mTOR, rather than the indirect influence of mTOR inhibition on cell size. To rule this out, we exploited the separate timing of cell growth and mTOR signaling. Treating cells with the mTOR inhibitor Torin-2, which inhibits cell growth, reduces cell size by over 15% (Figure 4A). However, when Torin-2 is washed out, mTOR activity is restored within 1 hour, whereas recovery of cell size proceeds over a period of 14 hours. This “size-recovery experiment” provides an opportunity to observe cells that are smaller than their regular size but are no longer subject to mTOR inhibition (Figure 4B). Western-blots of lysates from cells during the recovery phase show that the p38 MAPK remains upregulated after mTOR has resumed normal activity (Figure 4B). Further, the timescale of p38 dynamics during the recovery experiments, mirrors the recovery in cell size and not the dynamics of mTOR activity.

**Figure 4.**
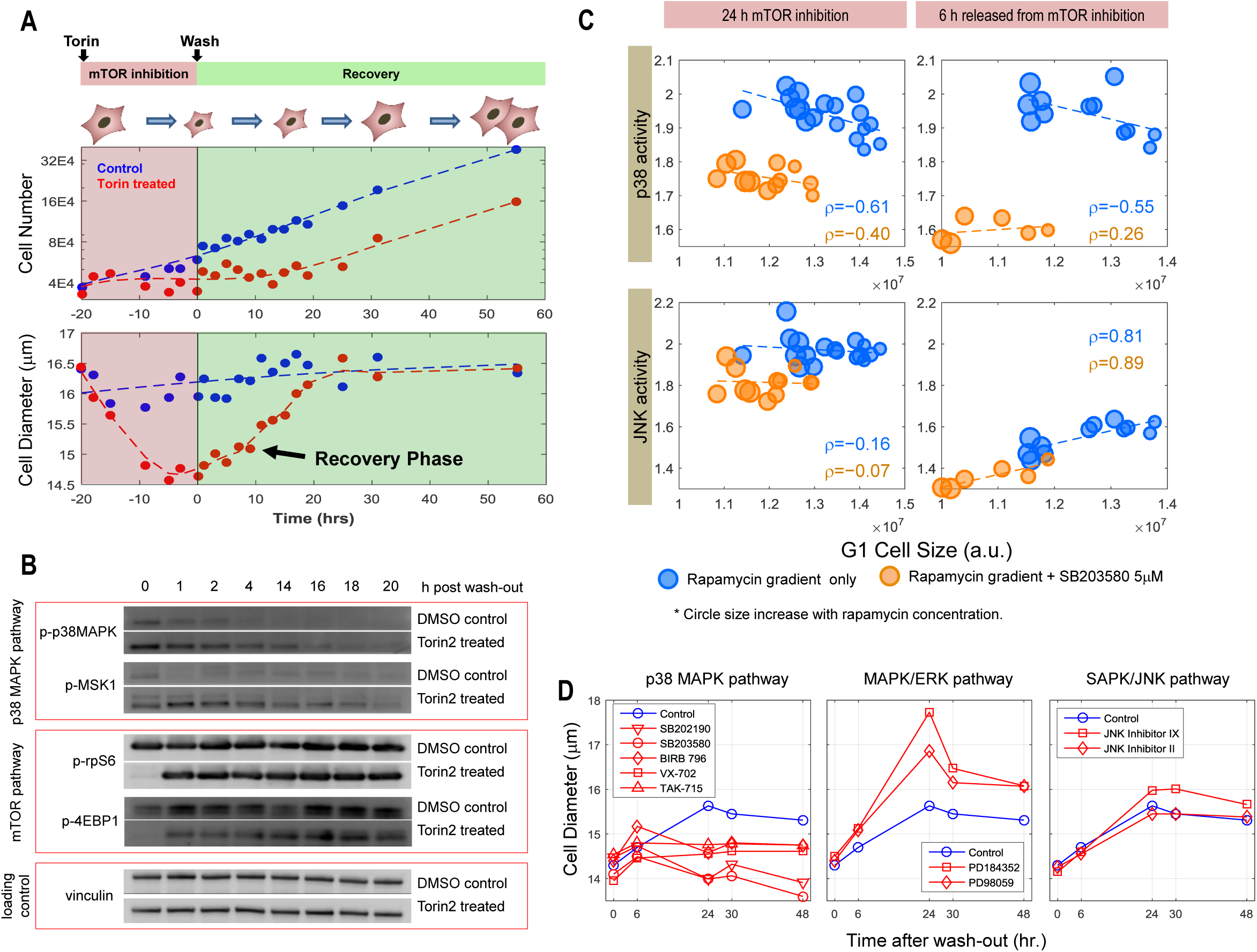
The p38 MAPK pathway is upregulated in small cells. **A**. Cells were treated with either 50 nM of Torin-2 or DMSO (control) for 20 hours, followed by drug wash-out and media replacement. Cells undergoing mTOR inhibition decrease in size and slow their proliferation rate. Following release from mTOR inhibition, cells recover in size while maintaining a low proliferation rate. Cells resume a wild type rate of proliferation only when their size reaches the size of the untreated population. **B**. Western-blots of whole cell lysates from 0 to 20 hours after release from mTOR inhibition. Level of mTOR pathway activity recovers within 1 hour after wash-out. The p38 MAPK is upregulated in the Torin-treated cells compared with controls, and the upregulation gradually fades away as cell size recovers. **C**. The p38 MAPK, but not JNK pathway is upregulated in small cells both under mTOR inhibition and after release from mTOR inhibition. Each data point is measured from a cell population with a minimum of 3,000 cells. The results shown here are representative of three independent experiments. **D**. Inhibition of p38 pathway, but not ERK or JNK pathway, represses recovery of size in cells released from mTOR inhibition. The cells were treated with 50 nM Torin-2 with or without (control) the indicated MAPK inhibitors for 22 hours. The cells were then released from Torin-2, but still subjected to the indicated MAPK inhibitor. All MAPK inhibitors were administrated at a concentration consistent with the highest corresponding concentration used in **Figure 2** and **Figure 2 -Figure supplement 2**. The results shown here are representative of two replicate experiments.

To further ask whether the p38 MAPK pathway is selectively upregulated in small cells, we employed a p38 Kinase Translocation Reporter (KTR) (23) that quantitatively reports p38 activation in live cells. This assay involves a fluorescent reporter that, once phosphorylated by p38 MAPK, shuttles from the nucleus to the cytoplasm. The ratio of cytoplasmic to nuclear (C/N) fluorescence represents a quantitative measure of p38 activation. As expected, cells treated with a p38 inhibitor display a lower C/N ratio than untreated cells (Figure 4C). Cells expressing the p38 KTR were treated with a range of concentrations of mTOR inhibitors, decreasing cell size to varying extents. The amount of p38 activity was measured in each condition. The results show that p38 activity is upregulated proportionally to the decrease in cell size (Figure 4C). When cells are released from mTOR inhibition, the negative correlation between cell size and p38 activity persists for at least 6 hours after release, long after normal mTOR activity has been restored. As an additional control, we expressed a dual color KTR in the cells, reporting the activity of both p38 and JNK. These measurements showed that while p38 activity is negatively correlated with cell size, JNK activity is not (Figure 4C). Together, these results indicate that aberrantly small cells upregulate the p38 MAPK pathway.

As a final test of a role for p38 in the maintenance of cell size, we asked whether its activity is necessary for recovery from the decrease in size caused by mTOR inhibition. We repeated the “size recovery experiment” shown in Figure 4A, in the presence or absence of several different p38 inhibitors. As additional controls, we also used inhibitors against two other MAPKs, ERK and JNK. Consistent with our expectation, inhibition of p38, but not of JNK or ERK, repressed the recovery in cell size (Figure 4D). Further, inhibition of p38 increased proliferation as compared to control (Figure 4 -Figure supplement 1), supporting the hypothesis that p38 activation prevents cell cycle progression of cells smaller than their normal size. Interestingly, the influence of p38 inhibition on the recovery in cell size persists long after the inhibitors are washed out (Figure 4 -Figure supplement 2). While Torin-2-treated cells begin to recover their size immediately after mTOR inhibition is relieved, cells that are co-treated with both Torin-2 and a p38 inhibitor display a marked delay in recovery (Figure 4 -Figure supplement 2). This delay may hint that the cell growth cycle depends on a commitment point that is cell-cycle-stage-dependent, analogous to the restriction point that regulates the cell division cycle (24).

In this study, we have shown that the p38 MAPK pathway is involved in coordinating cell cycle progression with cell size. Aberrantly small cells display higher p38 MAPK activity and longer duration of growth in the G1 phase of cell cycle. Inhibition of p38 MAPK leads to loss of the compensatory G1 length extension in small cells, resulting in faster proliferation, smaller cell size and increased heterogeneity in size. These results suggest a model whereby the p38 MAPK pathway functions downstream of a cell size sensing process and feeds information about cell size to regulators of cell cycle. Mechanisms by which the p38 MAPK pathway regulates cell cycle progression have been well-established in literature (4, 20–22). Here, we have shown that such p38- dependent cell cycle control is regulated by cell size. Previously, p38 MAPK has been shown be activated in response to osmotic shock that alters cell volume (6–8). The results reported here show that the p38 pathway is activated in cells with reduced cell mass, revealing that a common pathway responds to changes in both cell volume and mass. This finding is not trivial, since, unlike cell mass, cell volume is a labile phenotype that changes over rapid time scales (minutes) and is modulated by ion channels and transporters. In contrast, cell mass changes more slowly (over hours) and is modulated by protein translation and central carbon metabolism.

In addition to identifying the roles of p38 in communicating information on size, our study introduced a new strategy to assay chemical screens, particularly of quantitative physiological processes. Screens typically are designed to identify compounds that inhibit a given process leading to a phenotype. In this study we designed a strategy to identify compounds that perturb the relationship between two separate processes: growth and cell proliferation. We hope that strategies such as this may find further utility in drug discovery and other branches of cell physiology.

## Acknowledgments

We thank the Canadian Institutes of Health Research for its grant to R.K. (FRN-343437), and the National Institute of General Medical Sciences for its grant to M.K. (GM26875). We also thank Patricia and Alexander Younger and the Younger foundation for their generous donation to support this research. S.L. was supported by a graduate studentship award from the Research Training Center at the Hospital for Sick Children.

## Materials and Methods

### Materials

Succinimidyl ester conjugated to Alexa Fluor® 647 and DAPI were purchased from Life Technologies (Burlington, ON). The following small molecule inhibitors were purchased from Selleckchem (Houston TX): Rapamycin, Torin2, SB203580, VX-702, SB202190 (FHPI), BIRB 796 (Doramapimod), FR 180204, PD184352 (CI-1040), PD98059, JNK Inhibitor II, JNK Inhibitor IX. Lentiviral expression vectors encoding the JNK and p38 MAPK Kinase Translocation Reporters (KTR) were a kind gift from Markus Covert (Addgene plasmids No. 59151 and 59155). ON-TARGETplus SMARTpool siRNAs for the genes of interest as well non-targeting negative control siRNAs were obtained from Dharmacon (Lafayette, CO).

### Cell culture

HeLa (ATCC) and the retinal pigmented epithelial (RPE1, ATCC) cell lines stably expressing the degron of Geminin fused to Azami Green were cultured in DMEM medium (Life Technologies) supplemented with 10% Fetal Bovine Serum (FBS, Wisent, Montreal, QC) at 37 °C in a humidified atmosphere with 5% CO2. RPE1 cells stably co-expressing H2B conjugated to mTurquoise and Geminin conjugated mVenus were cultured in DMEM/F12 medium (Life Technologies) supplemented with 10% FBS. RPE1 cells stably expressing the JNK or p38 MAPK KTRs were generated as described below. Briefly, Human embryonic kidney HEK293T (ATCC) cells were maintained in DMEM (Wisent) supplemented with 10% FBS at 37°C and 5% CO2. The lentiviral transfer plasmids encoding the KTRs were co-transfected with plasmids encoding the packaging genes (viral Gag-pol) and the envelope gene, VSV-G, into the packaging HEK293T cells using jetPRIME transfection reagent (Polyplus Transfection New York, NY). The medium was changed 24h post-transfection and the viral supernatants were harvested 48h later, passed through a 0.45 μm syringe filter and frozen at -80 °C. The retrovirus was thawed at room temperature and then used to transduce RPE1 cells with the respective lentiviral transduction particles. Resistant clones were selected in 4 μg/mL puromycin (InvivoGen, San Diego, CA) for 3 days, isolated using cloning cylinders, and subsequently expanded and maintained in puromycin-containing DMEM medium. In all experiments, cell density was monitored to avoid over-confluence and contact inhibition.

### Compound screen

Two internal Novartis sets of tool compounds were screened, the publicly-known subset of compounds in the Mechanism-of-Action (MOA) Box (1,609 compounds) and a Kinome Box (1,637 compounds). The MOA Box is a dynamically-managed annotated list of compounds that cover target, pathway, and bioactivity space as comprehensively as possible to facilitate biological discovery by screening and profiling experiments. Bioactivity annotations were derived from integrated in-house assays and external assay data sources containing a mixture of qualitative target assignments (Thomson Reuters Integrity, DrugBank, Novartis-nominated MOA Box members) or quantitative dose response-type experimental assay data (chEMBL, GVK GOSTAR, Novartis assay data). Compounds were prioritized per target based on availability, amount of target evidence and, if available, clinical phase, while limiting the number of compounds selected from any one assay, publication, or patent. At the time of screening, chemical modulators of 903 unique human primary targets were represented based on hand-annotation, typically by 1-5 compounds/target where a compound-target association was made. In total, 42% of MOA Box compounds have only 1 assigned primary target, and a mean of 2 primary targets. However, additional target coverage for MOA Box compounds could be inferred from, the integrated sources listed above, comprising 1,964 total unique targets, such that 11% of members have only 1 inferable target and a median of 6 inferable targets. Target enrichment analysis, described below, was carried out using all possible targets (assigned and inferred from data). The target class coverage by percent of the MOA Box is as follows (**Figure 1 -Figure supplement 1**): Non-kinase Enzymes (28%), Kinases (16%), GPCR (15%), Proteases (12%), Ion Channel (11%), Unclassified (4%), Transporters (3%), Other Receptors (3%), Nuclear Hormone Receptors (3%), Cytokines (3%), and <1% each of Transcriptional Regulators, Signal Transducers and Structural Molecules. To assemble the Kinome Box, all physically-available inhibitors with publically-known structures having IC50 values <1 μM for a human kinase were filtered for only those inhibiting <25 total kinases in integrated internal and external data. A total of 473 human kinases were covered by 1,637 compounds (mean and median number of targets per compound equal to 5.4 and 2, respectively).

The screen was performed in 384-well μclear microplates (Greiner Bio-one, Monroe, NC). On day1, 2000 cells per well were seeded in 30 μl medium. On day2, compound was added to a final concentration of 1μM, 3 μM or 10μM (or 2μM, 6 μM or 20μM for Kinome Box) from a 2mM or 10mM stock solution using a Biomek FX with 384-well head (Beckman Coulter, Indianapolis, IN). The DMSO concentration was kept below or at 0.5% v/v. To reduce stochastic noise and promote overall screen accuracy, each compound was screened with 3 concentrations and duplicates.

### Compound treatment

The cells were seeded into 96-well μclear microplates (Greiner Bio-one, Monroe, NC) 16 hours prior to drug treatment. To perturb the p38, JNK and Erk pathways, the following specific inhibitors were used at the concentrations indicated in the figure legends. All compounds were dissolved in DMSO and diluted in DMEM when used. The concentrations have been selected to avoid severely interfering with cell proliferation or cause noticeable cell death. Twenty-four hours post compound treatment (control wells were treated with DMSO), the cells were treated and incubated with Rapamycin (30, 3, 1, and 0.3 nM) for an additional 24 hours before fixation and staining, or for whole cell lysis. DMSO only (0.01-0.5% v/v in DMEM) was used as a negative control for all the compound treatment experiments.

To examine the process of recovery in cell size, cells were treated with Torin2 (50 and 25 nM) for 24 hours, washed with PBS and re-incubated with fresh media for the times indicated. Cells were resuspended using trypsin and cell size and cell density was measured using the Multisizer 4 Coulter Counter (Beckman-Coulter, Mississauga, ON) or collected for whole cell lysis at time points indicated in the figures.

In the experiment using p38 or JNK KTRs, cells were seeded in 96 well plates and treated with Rapamycin (30, 3, 1, 0.3 nM and DMSO controls) for 24 hours. The cells were either fixed and stained or washed with PBS and replaced with fresh DMEM media for 6 hours before fixation and staining.

### siRNA transfection

RPE1-Geminin cells were seeded in 6-well plates at a density of 2 × 10^5^ cells/well. Twenty four hours post-seeding, the cells were transfected with SMARTpool siRNA (25 nM) using DharmaFECT 1 transfection reagent according to the manufacturer's instructions. Twenty hours post transfection, the cells were re-suspended using 0.05% trypsin-EDTA (Wisent) and re-seeded into 96-well μclear microplates (Greiner Bio-one, Monroe, NC) at a density of 5,000 cells/well. Six hours after re-seeding, cells were treated with Rapamycin (30, 3, 1 and 0.3 nM) or DMSO control for 24 hours before fixation and staining.

### Fixation, staining and imaging

Cells were fixed in 4% paraformaldehyde (Electron Microscopy Sciences, Hatfield, PA) for 10 minutes, followed by permeabilization in cold methanol at -20°C for 5 minutes. Cells were stained for cell size with 0.4 μg/mL Alexa Fluor 647 conjugated succinimidyl ester (SE) for 2 hours to nonspecifically label total protein content. The cells were then labeled for DNA with 1 μg/mL DAPI for 10 min. The cells were either imaged using an IN Cell Analyzer 2000 HCA system (GE Healthcare Life Sciences, Pittsburgh, PA) microscope at 10X magnification (compound screen) or using the Operetta High-Content Imaging System (Perkin Elmer, Woodbridge, ON) at 20X magnification (compound treatment and time-lapse experiments).

### Whole cell lysis and western blotting

To prepare whole cell lysates, cells were rinsed with ice-cold PBS and solubilized with RIPA Lysis Buffer (Boston Bio-Products, Boston MA) [50 mM Tris-HCl, 150 mM NaCl, 5 mM EDTA, 1 mM EGTA, 1% NP-40, 0.1% SDS and 0.5% sodium deoxycholate, pH 7.4] supplemented with protease and phosphatase inhibitor Cocktail (Thermo Scientific). Protein concentration was determined using the BCA protein assay (Thermo Scientific) and suspended with 4X Bolt® LDS Sample Buffer and 10X Bolt® Reducing Agent and heated for 10 min at 70 °C. Samples of equal protein were resolved by SDS-polyacrylamide gel electrophoresis and subjected to immunoblotting for proteins as indicated. The antibodies used for immunoblotting were all purchased from Cell Signaling Technology (Beverly, MA). All western-blot results in the figures have been reproduced in replicate experiments with cell lysates samples prepared in independent experiments.

### Image processing and cell segmentation

Automated image-processing pipelines have been developed using Matlab. The general scheme includes 3 steps: correction for background fluorescence and/or uneven illumination, cell segmentation, and feature quantification. Each step has been optimized according to microscopes, experimental design (*e.g*., fixed or live cell) and the features needed. The same image processing pipeline and parameters were applied identically to all images from the same experiment.

To correct for background fluorescence, a background image was constructed and applied per channel. There are two scenarios to construct a background image: either by averaging images containing few or no cells (in the compound screen); or by averaging the background region across all images collected (experiments with KTR cells, and in the time-lapse imaging). Uneven illumination was corrected for images from the compound screen in which the problem is obvious. A flat-field image has been constructed and applied for all channels. The flat-field image was constructed based on the fact that 2N peak in DNA distribution is invariant to cell positions in the image.

In the step of cell segmentation, nucleuses were first spotted through a nuclear channel (DNA or H2B), and segmented by seed-based watershed. When cell channel exists, the segmented nucleuses were further used as seeds to segment cells (SE channel) by watershed. The nuclear and cell border were detected with thresholds in corresponding channels, which were either automatically detected or manually selected based on need.

Following segmentation, each individual cell was processed to collect for its quantitative features, including total/average fluorescence per channel, nuclear size, *etc*.

### Cell cycle stages

Cells were first partitioned, according to the nuclear DNA level, into G1 (2N), S (2N-4N) and G2 (4N) phase. Progression in G1 phase was further divided, based on fluorescence of nuclear Geminin (and nuclear Cdt1 if available) into early G1 (low Geminin), late G1 (medium Geminin, high Cdt1), and G1/S transition (higher Geminin, medium to high Cdt1). The thresholds were automatically detected based on distribution of DNA, log(Geminin) and log(Cdt1).

### Analysis of the compound screen

The analysis was performed per plate separately, due to observed variability among plates. After image-processing of the compound screen datasets, every well captures approximately 4,000 cells. For each well, both cell size at early G1 stage (*S*) and proportions of cells in early G1 stage (*P*) were collected. The minimum volume ellipsoid (MVE) estimator (2, 3) was applied to pick a smallest volume ellipsoid that consists of around 50% of the total sample. The MVE estimator was a reliable tool to identify robust trend and detect outliers in multivariate data. The subsample of minimum volume ellipsoid was used to calculate the robust covariance matrix (rCov), from which correlation coefficient between *S* and *P* was calculated per plate. Further, we calculated the Mahalanobis distance from each data point (a well) to the center of the ellipsoid, and performed thresholding (5% significance) to the Mahalanobis distance to detect the outlier wells. Intuitively, the outlier wells display drastic distinctive phenotype in early G1 cell size and/or its G1-length compared with the control wells (and most drug treatments). Based on rCov, the principle vector of [*S, P*] was calculated and the angle of each outlier well with reference to the major principle vector was computed. The outlier angles were used to classify on/off-axis outliers: outliers with an angle smaller than 45 degree were classified as on-axis outliers, and otherwise off-axis outliers. The on/off-axis outliers were further filtered separately to lower false-positive rate. In the compound screen, each compound has been tested for at least 6 times (3 concentrations with duplicates); and compounds that have only been identified as outlier once among all treatments were excluded from further analysis. Next with the robust outliers, we performed the target enrichment analysis using hypergeometric test (Fisher’s exact test) for each known target of the outliers to identify the target proteins that are significantly highly represented in the outlier compounds.

To identify compound treatments that increase cell size variability, median absolute deviation (MAD), a robust measure of variability, of cell size was calculated per well. We observed that cell size MAD is linearly correlated with median cell size (*r* = 0.964). Accordingly, we normalize the MAD by median cell size to obtain a robust measure of cell-to-cell variability within cell populations. For each well, its normalized MAD is compared to that of the control wells in the same plate to calculate a MAD score. To select for compound treatments with perturbed size variability, a threshold in MAD score was calculated based on the distribution of MAD scores of all control wells (1% significance). The outliers with higher MAD scores compared with the threshold were filtered for repeatability and enriched for target proteins the same way as described above.

### Time-lapse microscopy

RPE1 cells with stable expression of H2B-mTurquoise and Geminin-mVenus were seeded in 96-well μclear microplates (Greiner Bio-one, Monroe, NC) and grown in the incubator for at least 6 hours prior to imaging. The cells were imaged with the Operetta High-Content Imaging System. During the imaging, the plate was incubated in the live cell chamber (37°C, 5% CO_2_) and grown in the same medium (DMEM/F12, 10% FBS) as described above. Widefield fluorescent images of H2B-mTurquoise and Geminin-mVenus were collected every 15 minutes at 20X magnification for 50 hours. Under this experimental setting, the microscope could support imaging of up to 4 wells every time. This experiment has been repeated three times, producing similar results as shown in **Figure 1F**.

### Automated lineage tracking

The live-cell images were first processed with the cell-segmentation pipeline to obtain single-cell features including cell position, fluorescent intensity and nuclear size. Subsequently, cells from each time point *T* were compared with the ones from *T*+1 to track for cell motion and division by searching for the globally optimal matches between neighboring time points (4). Parameters used for tracking were automatically calculated based on distribution of cell features, and confirmed in subsamples by eye. As output from tracking, each track starts with either appearance (first time point, or cell move into image field) or cell birth (with known mother and sister cell), and end with disappearance (last time point, or cell move out of image field) or cell division (with known daughter cells). After tracking, the individual cell tracks were further filtered to select for accurate tracks with the full cell cycle captured. Specifically, the algorithm selects for cell tracks that start with typical features of cell birth, end with typical features of mitosis and have relatively smooth fluorescence dynamics. Further, for each full cell cycle track, swift rise in Geminin level was detected to quantify early G1 duration. Initial nuclear size is estimated by averaging the first 9 data points (∼ 2.5 hours) after cell birth to decrease noise. Nuclear size measurements collected during mitosis were excluded from analysis, as H2B-mTurquoise does not accurately depict nuclear size during mitosis when nuclear envelope breaks down and chromosome condenses.

---

**Figure 1 -Figure supplement 1.**
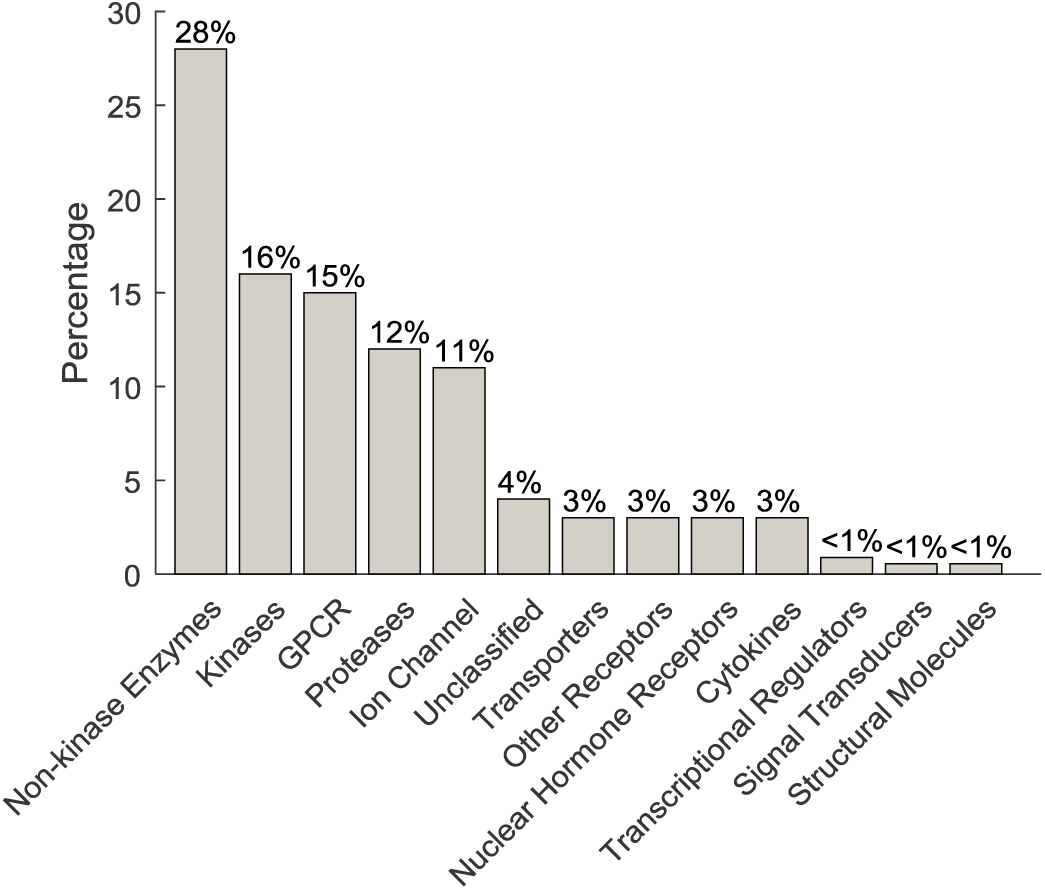
Percentage of the target class coverage of the MOA Box compounds.

**Figure 1 -Figure supplement 2.**
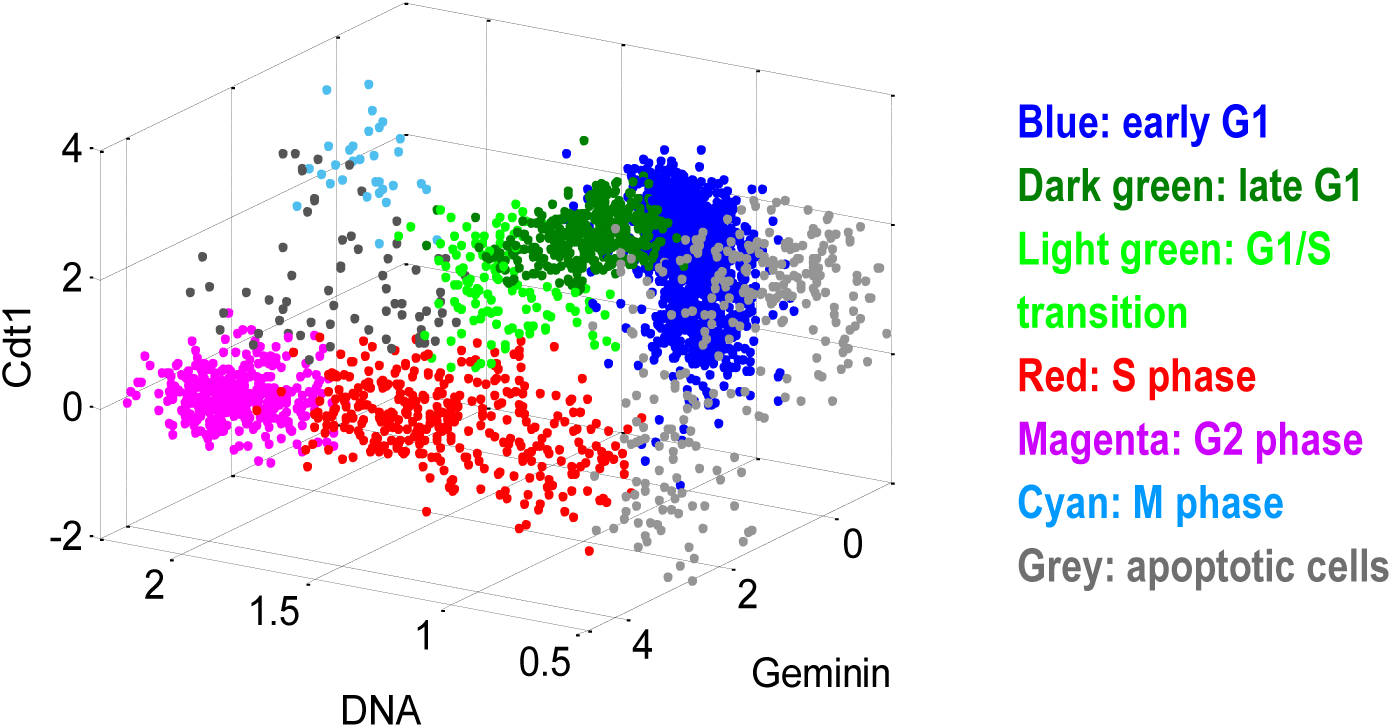
Cells from each well were partitioned, according to the 3 cell cycle indicators (DNA, Geminin, Cdt1), into discrete cell cycle stages. The scatterplot illustrates a control well (DMSO treated). Each point in the scatterplot is an individual cell. The DNA axis is normalized (1 = 2N, 2 = 4N). The Geminin and Cdt1 axes are in log scale (a.u.).

**Figure 1 -Figure supplement 3.**
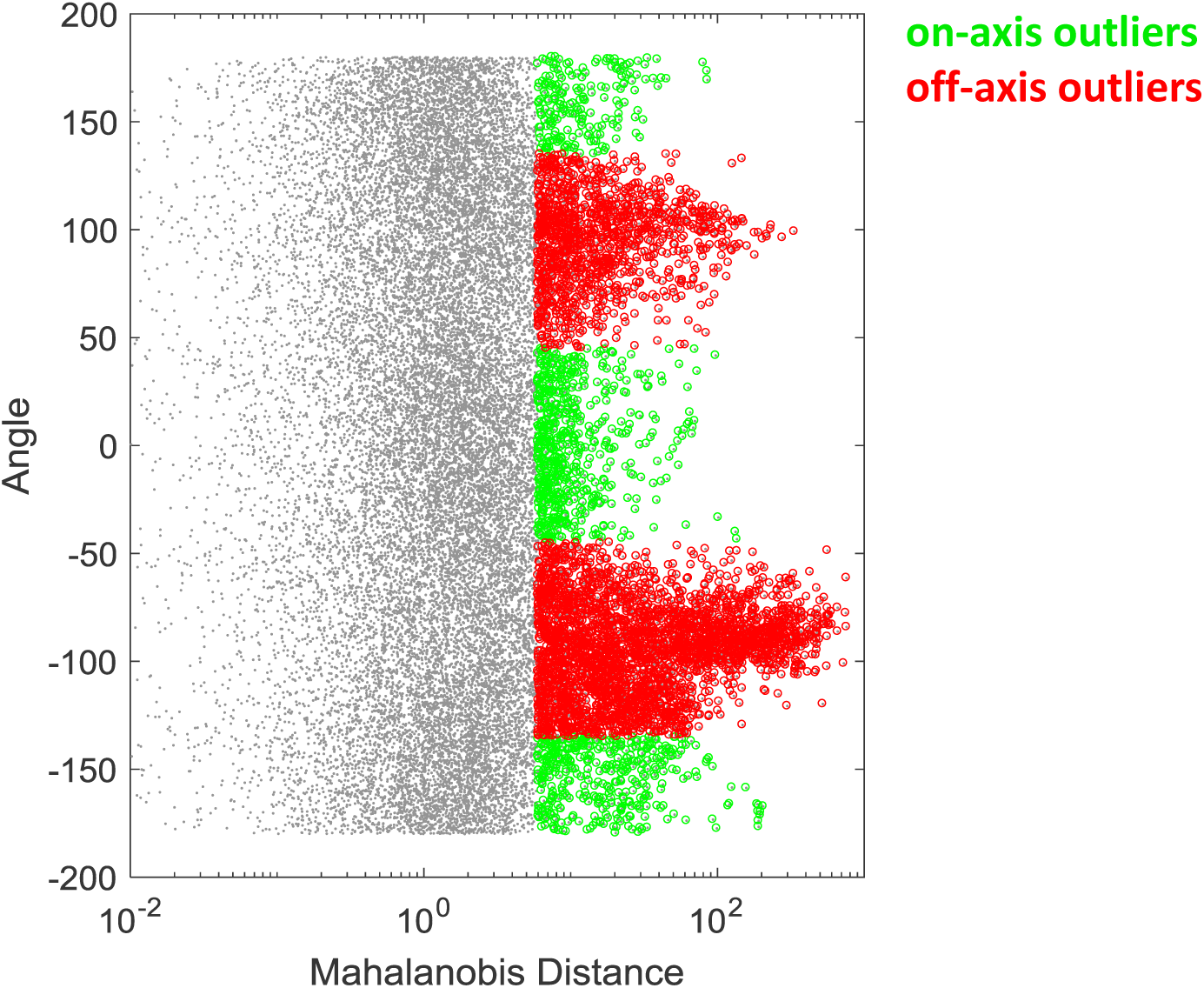
The outliers of the screen (as exemplified in **Figure 1B**) were identified by thresholding (5% significance) the Mahalanobis distance. The outliers were then classified into on-axis or off-axis based on the angle of the data points with reference to the major principle vector. Specifically, outliers with an angle smaller than 45 degree were classified as on-axis outliers, and otherwise off-axis outliers. See **Material and Methods -Analysis of the compound screen** for details.

**Figure 1 -Figure supplement 4.**
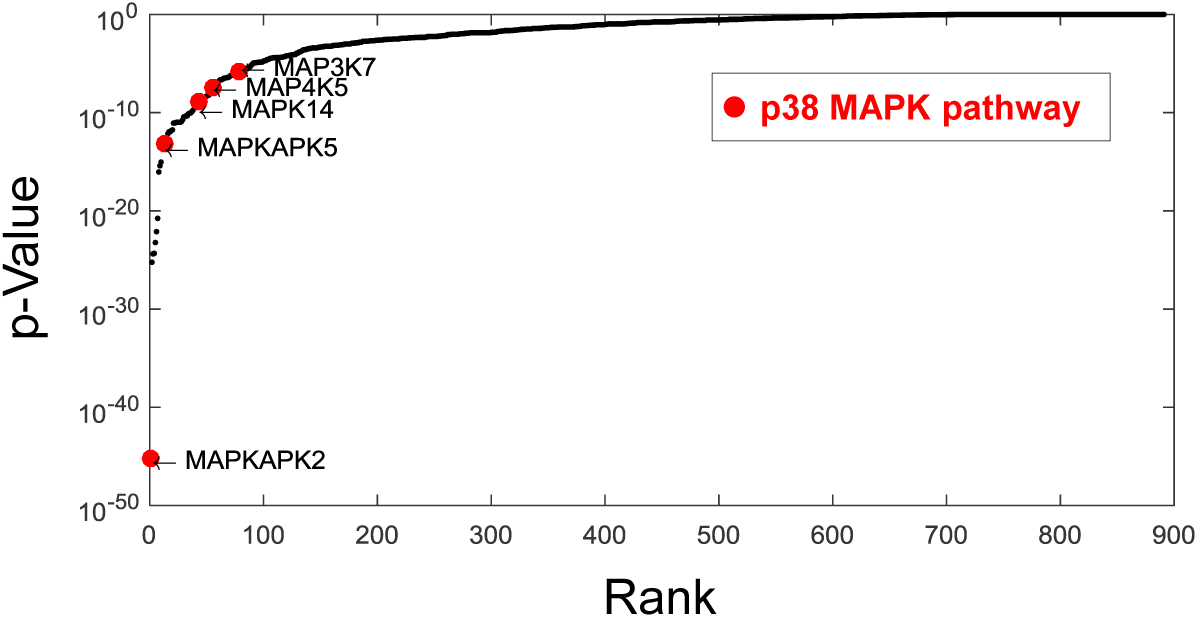
Ranked *p*-values from the enrichment analysis of compounds that increase cell size variability (Fisher’s exact test). Components from the p38 pathway (highlighted) were highly enriched. Specifically, MK2/MAPKAPK2, a direct downstream substrate of p38 is the top-ranking genes that associate with increased cell size variability.

**Figure 2 -Figure supplement 1.**
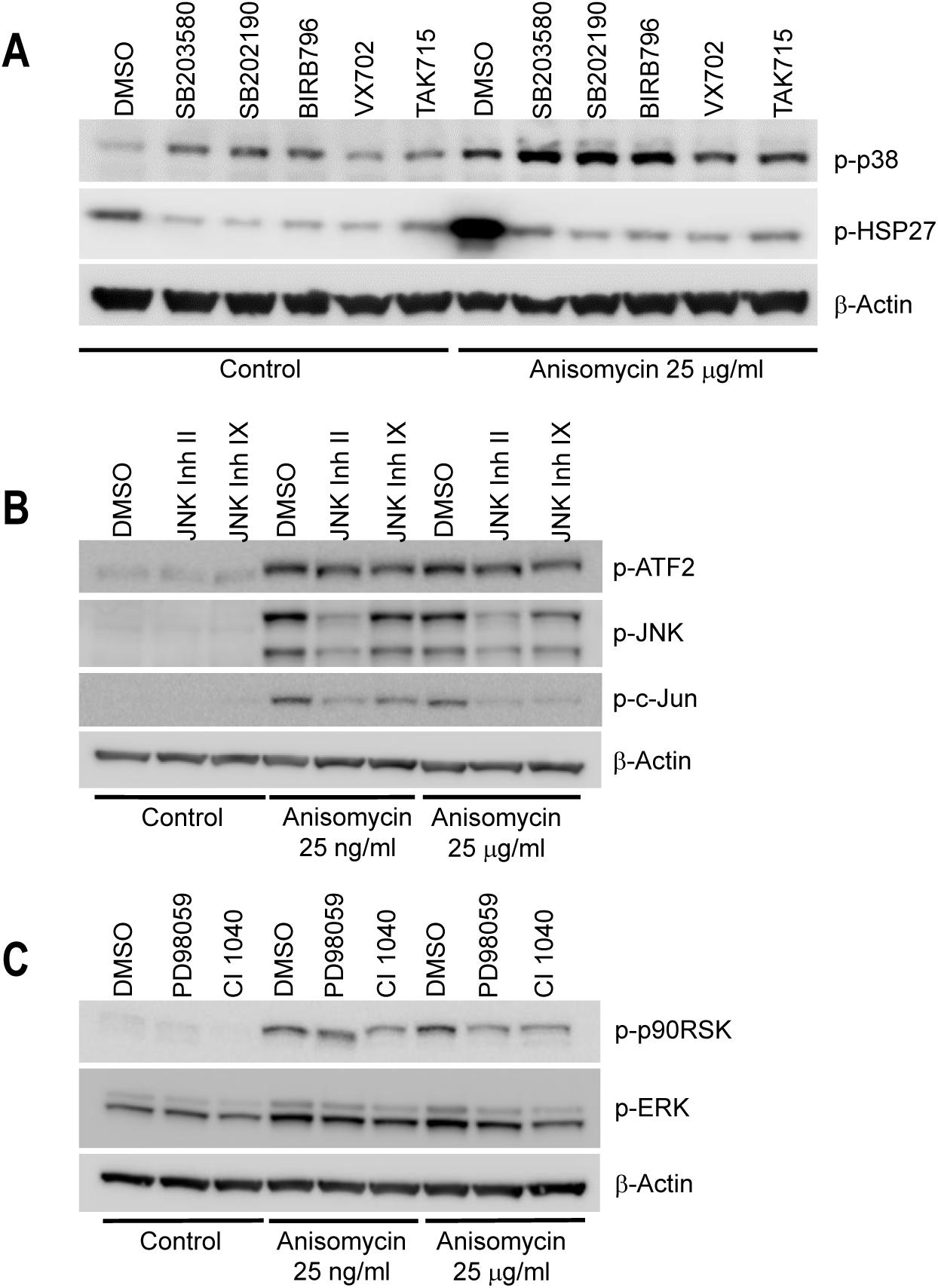
Western-blot of cell lysates from conditions shown in **Figure 2C** confirms the chemical inhibitors are efficient towards inhibiting corresponding MAPKs pathway. Cells were treated with indicated inhibitors for 24 hours before collecting lysates. Anisomycin was added to select wells 1 hour prior to making lysates, to activate MAPK pathways. All inhibitors were used at the ‘high dose’ indicated in **Figure 2C** and **Figure 2 -Figure supplement 2**. **A.** Cells treated with p38 inhibitors display a lower level of p-HSP27 (downstream of p38). The p38 inhibitors induce a higher level of p-p38. This is due to negative feedback in the p-p38 pathway (1), and the fact that p38 inhibitors prevent p-p38 from phosphorylating downstream substrates, but do not block phosphorylation of p38 itself by upstream regulators. **B, C.** Cells treated with JNK or MEK I/II inhibitor inactivate the corresponding pathway under Anisomycin induction. The influence of the inhibitor is not obvious under control condition probably due to low basal activation of the pathways.

**Figure 2 -Figure supplement 2.**
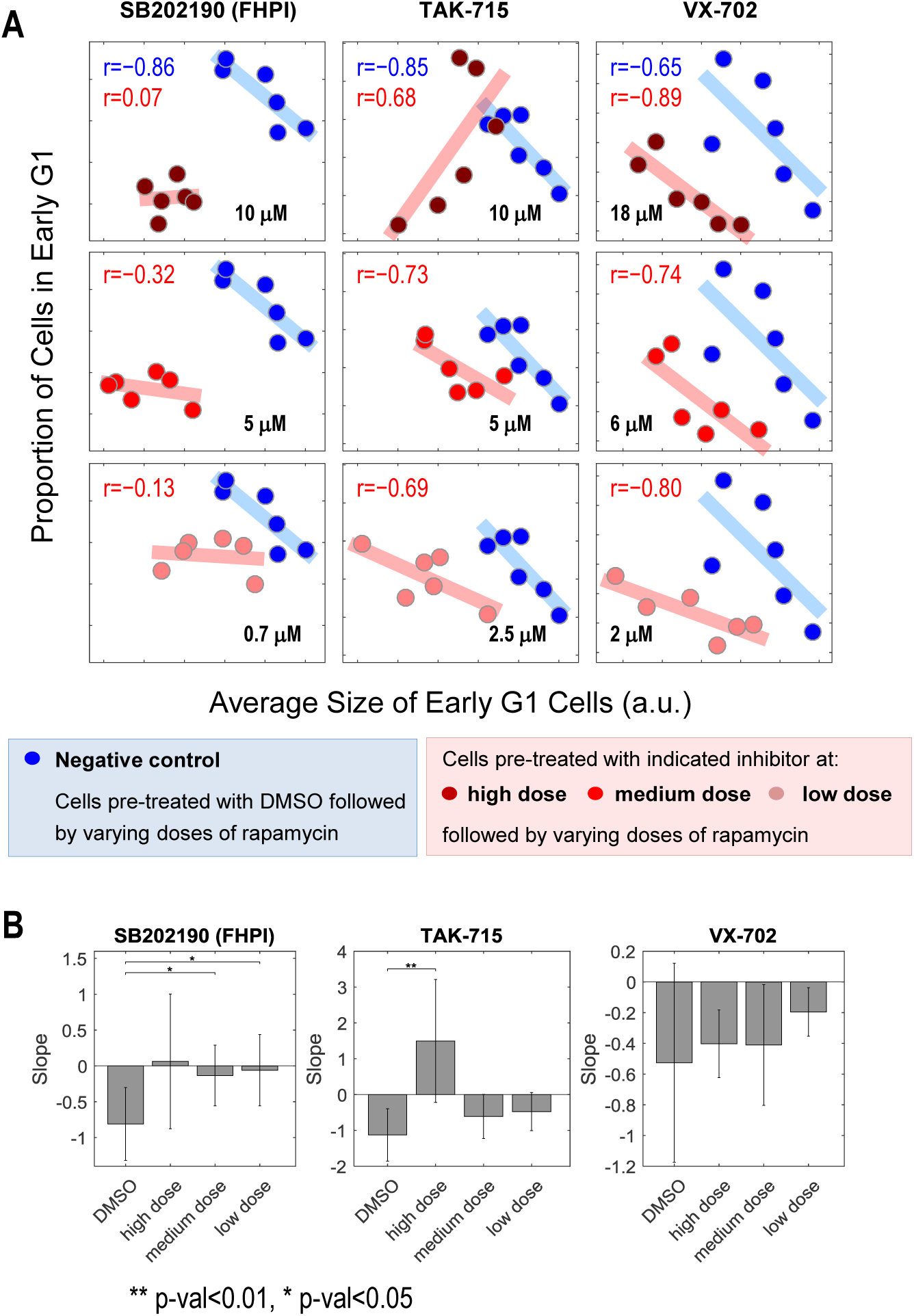
The negative correlation between cell size and proportion of cells in early G1 is perturbed or weakened under p38 inhibition. Measurements collected in the same experiment as **Figure 2C**. **A**. Scatterplot comparing cells of negative control (DMSO) with cells under p38 inhibition (treated with indicated inhibitor and concentration). Each data point was measured from a cell population with a minimum of 7,000 cells. The results are representative of three independent experiments. **B**. The slope between size and proportion of cells in G1 is either disturbed or weakened. *p*-values were calculated with one-tailed Student *t*-test (H_0_: slope of control>=slope of compound treatment).

**Figure 3 -Figure supplement 1.**
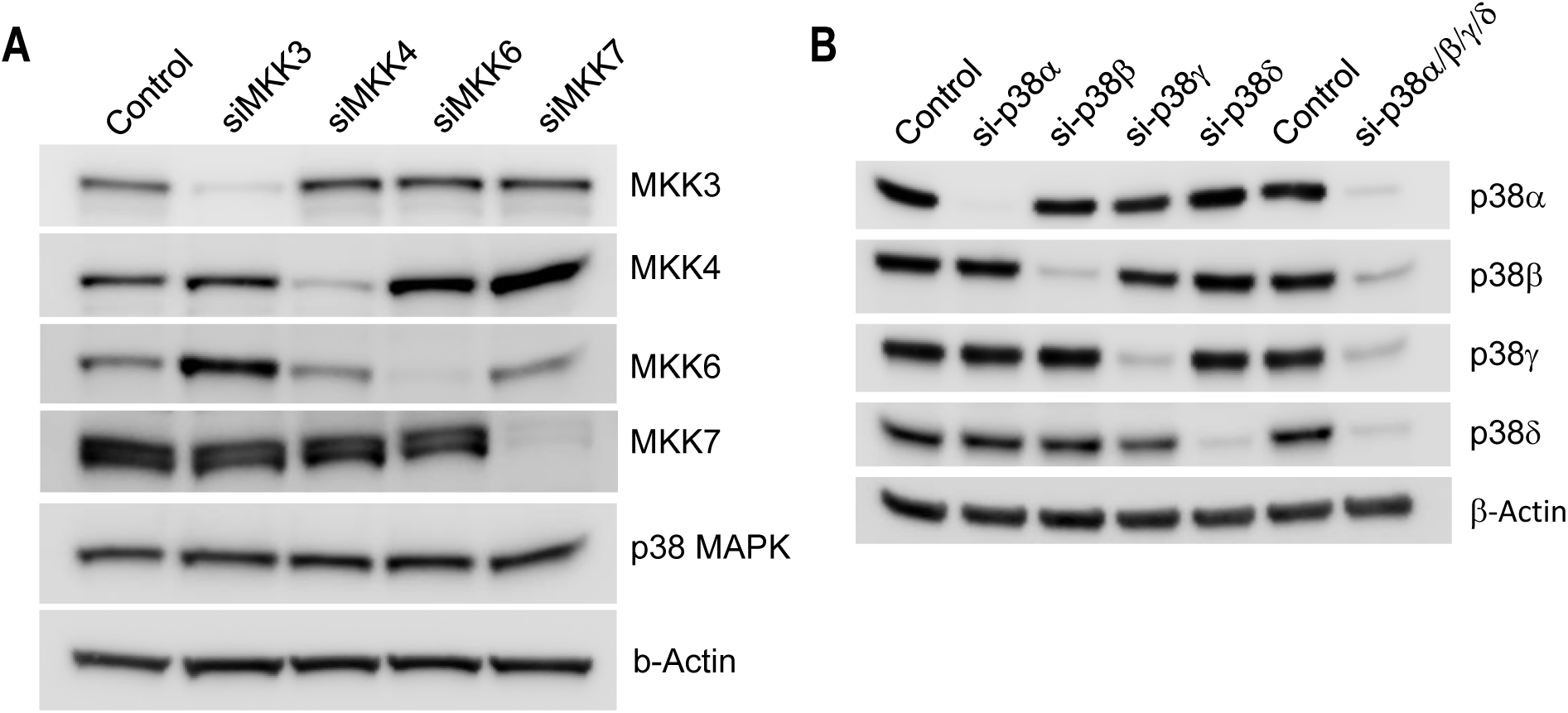
Western-blot of cell lysates from conditions shown in Fig 3 confirms efficiency of knockdown of MKKs (**A**) or p38 isoforms (**B**).

**Figure 4 -Figure supplement 1.**
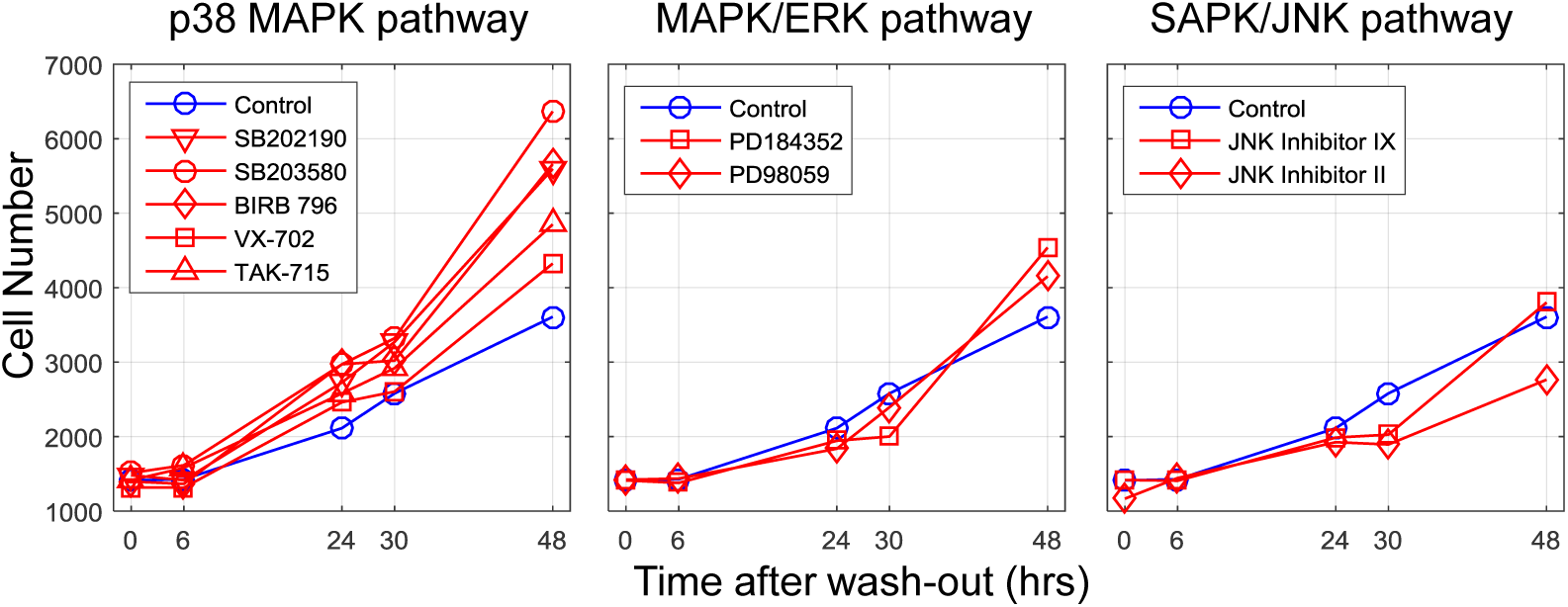
Cells under inhibition of p38, but not Erk or JNK pathway, increased cell proliferation compared with control conditions when cells were released from mTOR inhibition. The cells were co-treated with Torin-2 (50 nM) together with the indicated MAPK inhibitor for 22 hours, and then released from Torin-2 but still subjected to the indicated MAPK inhibitor. The data were collected from the same experiment as **Figure 4D**. The results shown here are representative of two replicate experiments.

**Figure 4 -Figure supplement 2.**
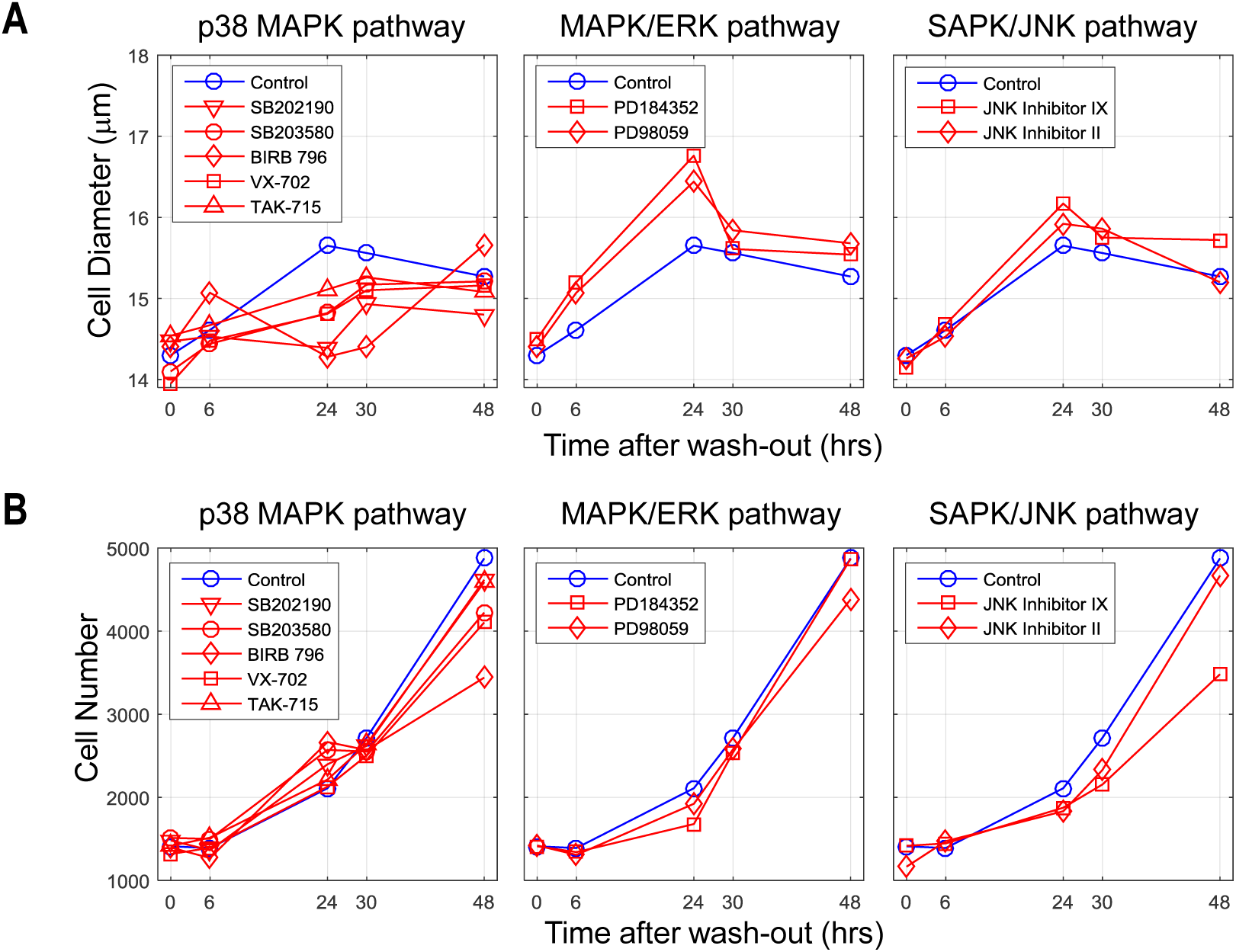
Recovery in cell size is delayed even after the p38 inhibitor was wash-out. Cells were co-treated with both Torin-2 (50 nM) and the indicated MAPK inhibitor for 22 hours. The cells were then released from both inhibitors. Both cell size (**A**) and cell number (**B**) were measured. The results shown here are representative of two replicate experiments.

## References

1. M. B. Ginzberg, R. Kafri, M. Kirschner, On being the right (cell) size. Science. 348, 1245075 (2015).

2. I. Rupes, Checking cell size in yeast. Trends Genet. TIG. 18, 479–485 (2002).

3. A. Sveiczer, B. Novak, J. M. Mitchison, The size control of fission yeast revisited. J. Cell Sci. 109 (Pt 12), 2947–2957 (1996).

4. T. M. Thornton, M. Rincon, Non-Classical P38 Map Kinase Functions: Cell Cycle Checkpoints and Survival. Int. J. Biol. Sci. 5, 44–52 (2008).

5. C. Ambrosino, A. R. Nebreda, Cell cycle regulation by p38 MAP kinases. Biol. Cell. 93, 47–51 (2001).

6. J. Han, J. D. Lee, L. Bibbs, R. J. Ulevitch, A MAP kinase targeted by endotoxin and hyperosmolarity in mammalian cells. Science. 265, 808–811 (1994).

7. T. Moriguchi et al., Purification and identification of a major activator for p38 from osmotically shocked cells. Activation of mitogen-activated protein kinase kinase 6 by osmotic shock, tumor necrosis factor-alpha, and H2O2. J. Biol. Chem. 271, 26981–26988 (1996).

8. L. New, J. Han, The p38 MAP Kinase Pathway and Its Biological Function. Trends Cardiovasc. Med. 8, 220–228 (1998).

9. A. Clerk, A. Michael, P. H. Sugden, Stimulation of the p38 mitogen-activated protein kinase pathway in neonatal rat ventricular myocytes by the G protein-coupled receptor agonists, endothelin-1 and phenylephrine: a role in cardiac myocyte hypertrophy? J. Cell Biol. 142, 523–535 (1998).

10. S. Kudoh et al., Mechanical Stretch Induces Hypertrophic Responses in Cardiac Myocytes of Angiotensin II Type 1a Receptor Knockout Mice. J. Biol. Chem. 273, 24037–24043 (1998).

11. Á. Molnár, A. M. Theodoras, L. I. Zon, J. M. Kyriakis, Cdc42Hs, but Not Rac1, Inhibits Serum-stimulated Cell Cycle Progression at G1/S through a Mechanism Requiring p38/RK. J. Biol. Chem. 272, 13229–13235 (1997).

12. S. López-Avilés et al., Inactivation of the Cdc25 Phosphatase by the Stress-Activated Srk1 Kinase in Fission Yeast. Mol. Cell. 17, 49–59 (2005).

13. M. Cully et al., A role for p38 stress-activated protein kinase in regulation of cell growth via TORC1. Mol. Cell. Biol. 30, 481–495 (2010).

14. A. Sakaue-Sawano et al., Visualizing spatiotemporal dynamics of multicellular cell-cycle progression. Cell. 132, 487–498 (2008).

15. R. Kafri et al., Dynamics extracted from fixed cells reveal feedback linking cell growth to cell cycle. Nature. 494, 480–483 (2013).

16. B. González-Terán et al., P38γ and δ promote heart hypertrophy by targeting the mTOR-inhibitory protein DEPTOR for degradation. Nat. Commun. 7, 10477 (2016).

17. B. Snijder et al., Single-cell analysis of population context advances RNAi screening at multiple levels. Mol. Syst. Biol. 8, 579 (2012).

18. J. T. Marques, B. R. G. Williams, Activation of the mammalian immune system by siRNAs. Nat. Biotechnol. 23, 1399–1405 (2005).

19. R. Grumont et al., The mitogen-induced increase in T cell size involves PKC and NFAT activation of Rel/NF-kappaB-dependent c-myc expression. Immunity. 21, 19–30 (2004).

20. J. N. Lavoie, G. L’Allemain, A. Brunet, R. Müller, J. Pouysségur, Cyclin D1 Expression Is Regulated Positively by the p42/p44MAPK and Negatively by the p38/HOGMAPK Pathway. J. Biol. Chem. 271, 20608–20616 (1996).

21. K. Mikule et al., Loss of centrosome integrity induces p38—p53—p21-dependent G1—S arrest. Nat. Cell Biol. 9, 160–170 (2007).

22. A. S. Yee et al., The HBP1 transcriptional repressor and the p38 MAP kinase: unlikely partners in G1 regulation and tumor suppression. Gene. 336, 1–13 (2004).

23. S. Regot, J. J. Hughey, B. T. Bajar, S. Carrasco, M. W. Covert, High-Sensitivity Measurements of Multiple Kinase Activities in Live Single Cells. Cell. 157, 1724–1734 (2014).

24. M. V. Blagosklonny, A. B. Pardee, The restriction point of the cell cycle. Cell Cycle Georget. Tex. 1, 103–110 (2002).

## References

1. J. S. C. Arthur, S. C. Ley, Mitogen-activated protein kinases in innate immunity. Nat. Rev. Immunol. 13, 679–692 (2013).

2. P. J. Rousseeuw, B. C. van Zomeren, Unmasking Multivariate Outliers and Leverage Points. J. Am. Stat. Assoc. 85, 633–639 (1990).

3. P. J. Rousseeuw, Multivariate estimation with high breakdown point. Math. Stat. Appl. 8, 283–297 (1985).

4. T. Kanade et al., in 2011 IEEE Workshop on Applications of Computer Vision (WACV) (2011), pp. 374–381.

